# A high-carbohydrate diet prolongs dysbiosis and *Clostridioides difficile* carriage and increases late-onset CDI and mortality in a hamster model of infection

**DOI:** 10.1101/2021.10.04.463142

**Authors:** Shrikant S. Bhute, Chrisabelle C. Mefferd, Jacqueline R. Phan, Muneeba Ahmed, Amelia E. Fox-King, Stephanie Alarcia, Jacob V. Villarama, Ernesto Abel-Santos, Brian P. Hedlund

## Abstract

Studies using mouse models of *Clostridioides difficile* infection (CDI) have demonstrated a variety of relationships between dietary macronutrients on antibiotic-associated CDI; however, few of these effects have been examined in hamster models of CDI. In this study, we investigated the effect of a high-carbohydrate diet previously shown to protect mice from CDI on the progression and resolution of CDI in a hamster disease model. Hamsters fed the high-carbohydrate diet developed distinct diet-specific microbiomes during antibiotic treatment and CDI, with lower diversity, persistent *C. difficile* carriage, and delayed microbiome restoration. In contrast to 0% mortality in mice, 80% of hamsters fed the high-carbohydrate diet developed fulminant CDI and died, including several cases of late-onset CDI, whereas only 33% of hamsters fed a standard lab diet developed CDI only during the acute phase. We speculate that prolonged dysbiosis in these animals allowed *C. difficile* to proliferate following a three-day vancomycin course administered as part of this model system, leading to secondary CDI and eventual mortality. This study, along with similar studies in mouse models of CDI, suggests high-carbohydrate diets promote antibiotic-associated dysbiosis and long-term *C. difficile* carriage, which may convert to symptomatic CDI when conditions change.

**Importance:** The effects of diet on CDI are not completely known, although most studies in mouse CDI models show that dietary carbohydrates ameliorate CDI. Here, we used a high-carbohydrate diet previously shown to protect mice against CDI to assess its effect on a hamster model of CDI and paradoxically found that it promoted dysbiosis, *C. difficile* carriage, and higher mortality. A common thread in both mouse and hamster experimental models was that the high-carbohydrate diet promoted long-term carriage of *C. difficile*, which may have converted to fulminant CDI only in the highly susceptible hamster model system. If diets high in carbohydrates also promote dysbiosis and *C. difficile* carriage in humans, then these diets might paradoxically increase chances of CDI relapse despite their protective effects against primary CDI.

## Introduction

*Clostridioides difficile* infection (CDI) is caused by toxigenic strains of the anaerobic, Gram-positive, spore-forming enteropathogen *Clostridioides* (formerly *Clostridium*) *difficile*. CDI is the most commonly identifiable cause of antibiotic-associated diarrhea. The incidence of CDI is complicated by the appearance of highly resistant and hypervirulent BI/NAP1/027 strains that have an attributable mortality of an astonishing 22% (1). In the US, hospital- and community-acquired CDI accounted for 223,900 cases with 12,800 deaths in 2017 (2), leading to a significant financial burden ($4.8 billion) on the US healthcare system (2–4).

The outcome of CDI varies widely from asymptomatic colonization to mild and self-limiting diarrhea, to severe diarrhea and life-threatening pseudomembranous colitis (5, 6). The diverse and unpredictable CDI outcomes can be linked to both *C. difficile*-specific and host-related factors. *C. difficile* pathogenicity loci vary considerably from strain to strain (7, 8), leading to the variable presence of pathogenesis factors. For example, the presence of the two large clostridial toxins (TcdA and TcdB) is strain-dependent and can be attributed to different CDI outcomes (9). *C. difficile* is also catabolically versatile and is able to use both diet- and host-derived amino acids and sugars as resources depending on diet or during the course of infection (10). In addition, host-derived primary bile acids act as spore co-germinants that can amplify *C. difficile* growth in the gut (11). Yet, under normal circumstances, the normal gut flora provides colonization resistance to acute *C. difficile* infections (12), likely due to niche exclusion and the production of inhibitory volatile fatty acids and secondary bile acids (13). When the microbiota is disturbed (e.g., antibiotic treatment) *C. difficile* spores can germinate and establish infection.

Although diet is known to modulate gut microbial ecology (14, 15), studies on the effects of dietary macronutrients on CDI outcome have been largely restricted to small animal models, mostly mice (16–18). Recent studies conducted using mouse models of antibiotic-associated CDI have shown that high-protein diets lead to fulminant CDI and high mortality rates (16–18). These studies indicate some degree of consensus on the effect of protein-rich diets leading to severe CDI, but the effects of dietary carbohydrates appear to be more complex. For example, diets high in microbiota-accessible carbohydrates (MACs) that are fermented to short-chain fatty acids in the intestinal tract have been reported to be protective against CDI (17). Conversely, other studies have implicated simple carbohydrates, specifically trehalose, in the proliferation of hypervirulent *C. difficile* strains (19) and dietary glucose and fructose in enhancement of both spore production and host colonization capacity (20). In contrast, our previous study showed that a diet rich in carbohydrates decreased CDI severity and eliminated mortality in mice, despite the fact that the carbohydrates provided in the diet were readily digestible sucrose and starch (21). Interestingly, our study also indicated that the high-carbohydrate diet led to an asymptomatic persistent carrier state in which *C. difficile* was present in the murine gut for at least a month after primary CDI was resolved, with constant shedding into feces.

Our previous study and other similar studies have been carried out using a recently developed mouse model of CDI (22, 23), which is more resistant to the development of CDI over the more traditional, highly sensitive hamster models (24). For over five decades, Syrian golden hamster models of CDI have been extensively used to assess the growth dynamics and pathophysiology of *C. difficile*, the susceptibility and severity of CDI following different antibiotic regimens, and the efficacy of various treatment options (24–27). Hamsters are exquisitely sensitive to CDI and develop fulminant colitis within 48 hours post-challenge. These extreme symptoms are rarely seen in primary human CDI. However, hamster CDI severity can be modulated by vancomycin treatment, which is also used clinically to treat CDI. Indeed, suboptimal vancomycin treatment of infected hamsters results in delayed sign onset, similar to human CDI relapse, and also prevents non-CDI clindamycin-associated colitis (28). Previously, dietary supplementation of fructooligosaccharides (FOS) and soy fiber have been shown to delay CDI onset, attenuate CDI development, and increase survival time in hamsters (29, 30). Additionally, increased CDI susceptibility has been observed in hamsters fed an atherogenic diet (31). Despite these efforts, the effects of dietary macronutrients on CDI in hamster models of disease are not completely understood. Considering the lack of clinically translatable human studies indicating the precise effect of dietary macronutrients on CDI outcome, and the lack of consensus on the effects of carbohydrates on CDI in murine models (17–20), a better understanding of the effects of dietary carbohydrates on hamster models of CDI is likely to provide useful insights into *C. difficile* pathogenesis.

Here, we used a Syrian golden hamster model of CDI with a three-day vancomycin treatment to modulate CDI to assess the effect of a high-carbohydrate diet previously shown to be protective in mice (18) on the outcome of antibiotic-induced CDI using the toxigenic hypervirulent *C. difficile* strain R20291. The high-carbohydrate diet promoted antibiotic- and CDI-associated dysbiosis and *C. difficile* carriage, eventually leading to higher mortality following a subclinical vancomycin treatment used to modulate disease severity. Thus, although some high-carbohydrate diets can prevent severe CDI in some settings, they may also promote *C. difficile* carriage, which could change from an asymptomatic carrier state to severe CDI relapse depending on environmental cues.

## Materials and Methods

### Materials

Brain heart infusion (BHI) medium was purchased from BD Biosciences (Franklin Lakes, NJ). Reagents for DNA isolation were obtained from Qiagen (product no. 51504; Qiagen, Germantown, MD). Reagents for PCR were obtained from Quantabio (product no. 2200410; Quantabio, Beverly, MA).

### *C. difficile* growth conditions and spore harvest

*C. difficile* strain R20291 (RT027) was the kind gift of Dr. Nigel Minton (University of Nottingham). *C. difficile* cells were allowed to sporulate by growing at 37°C for 7 days on Brain Heart Infusion agar plates (Bacto) supplemented with 2% yeast extract, 0.1% L-cysteine HCl, and 0.05% sodium taurocholate. Plates were incubated in a Coy vinyl anaerobic chamber (Coy Lab Products, MI) containing 10% CO_2_, 10% H_2_, and 80% N_2_. Bacterial cells and spores were harvested from plates by washing with ice-cold, autoclaved, deionized water and gentle scraping. Vegetative cells and spores were pelleted by centrifugation and resuspended in fresh deionized water for a total of three wash cycles. Pellets were then passed through a HistoDenz gradient (20% −50%) to purify spores from cell debris and washed with autoclaved deionized water five times. Schaeffer-Fulton-staining was performed to ensure >95% purity of the harvested spores and the resulting purified spores were stored at 4°C until administration.

### Animals

Female Syrian golden hamsters were purchased from Charles River (Wilmington, MA) and acclimated for a week in the animal facility prior to the experiments. Autoclaved bedding, water, and feed were used in all procedures. The Institutional Animal Care and Use Committee (IACUC) at the University of Nevada, Las Vegas, reviewed and approved this study (R0914-297). All experiments were performed according to the National Institutes of Health guidelines in the Guide for Care and Use of Laboratory Animals. Experimental animals were 5-8 weeks old.

### Treatment groups, induction and monitoring of CDI, and sample collection

A total of 30 hamsters were housed in individual cages and fed a standard lab diet for 6 days (until Day −14). Hamsters were then randomly separated into three groups of 10 hamsters; two groups were fed a standard lab diet (Standard Lab Diet and Standard Lab Diet (-CDI)), and one group received a high-carbohydrate diet from Day −14 onward for the remainder of the experiment (High-Carbohydrate Diet) (**Figure 1**). Irradiated hamster chow was purchased from TestDiet and stored at 4°C before use (**Supplementary file 1 and 2**). These are the exact same diets used previously for experiments in a mouse model of CDI (18). Diets and autoclaved water were given *ad libitum*. The hamsters in the High-Carbohydrate Diet and Standard Lab Diet groups were given a single dose of clindamycin dissolved in autoclaved DI water (30 mg/kg) via oral gavage on Day −1, challenged with 10^2^ CFUs of *C. difficile* R20291 spores by oral gavage on Day 0, and given vancomycin dissolved in autoclaved DI water (1 mg/mL) via oral gavage on Days 0-2. Vancomycin was used to prevent non-CDI clindamycin-associated colitis (28), which led to death of animals before *C. difficile* challenge in pilot experiments, and to modulate CDI severity (21). The hamsters in the Standard Lab Diet (-CDI) group did not receive any antibiotic treatment and were not infected with *C. difficile* spores. All hamsters were monitored daily for signs of CDI for 14 days post-infection, weighed twice daily after challenge inside a biosafety cabinet, and euthanized 14 days post infection, or as soon as hamsters showed CDI signs (diarrhea/increase in soiled bedding), so that no animals experienced unrelieved pain or distress. Fecal samples were collected beginning at Day −20 and for the duration of the 34-day experiment as indicated in **Figure 1**, and archived for microbiome analysis at −80 °C.

**Fig 1.**
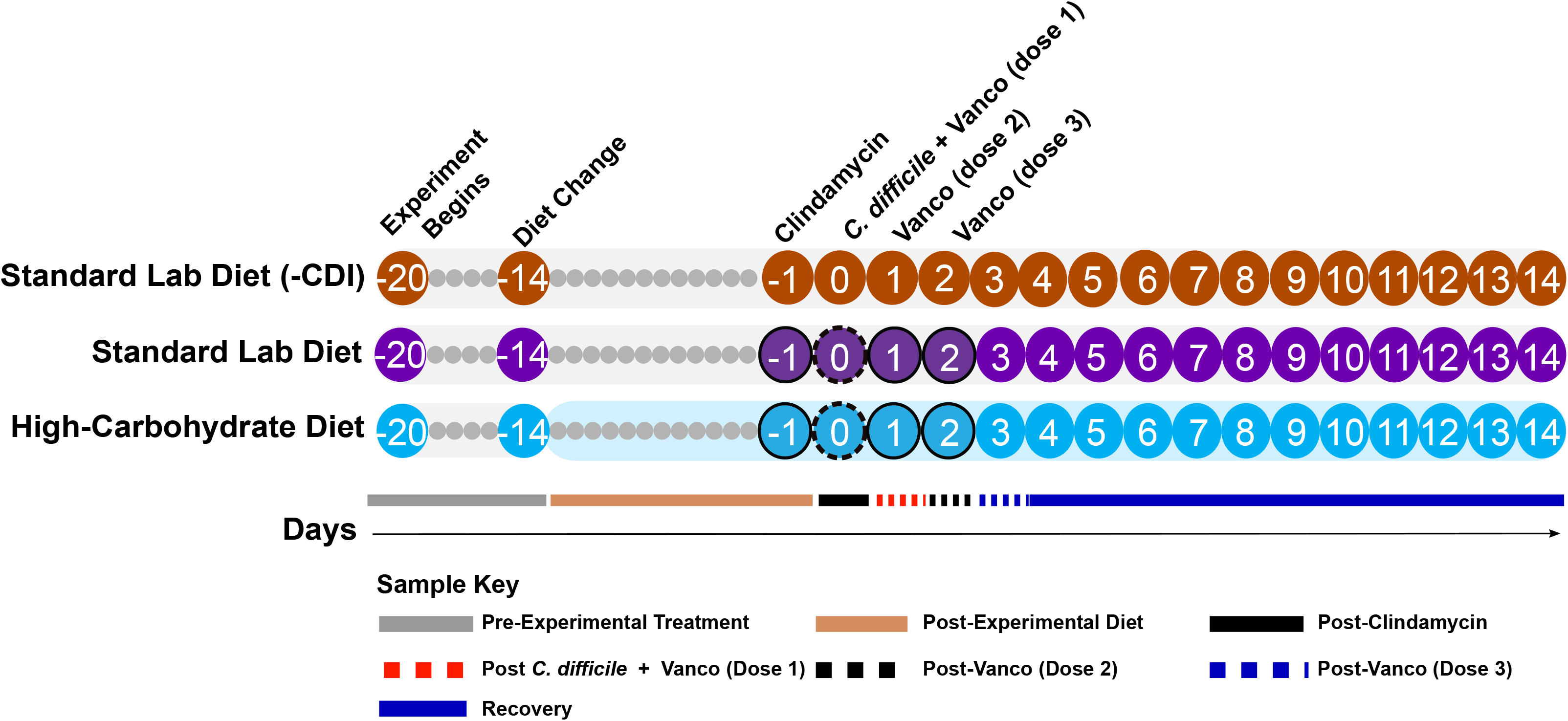
Experimental timeline and stool collection. The high-carbohydrate diet was introduced on Day 6 (−14). Clindamycin was administered on Day −1 to induce dysbiosis, and hamsters were challenged with *C. difficile* R2027 spores on Day 0. A vancomycin regimen (outline circle) to mitigate CDI was carried out on Days 0-2. Circles with numbers indicate days fecal samples were collected. Stool collection took place before the manipulation of hamsters or experimental treatment.

### DNA extraction and 16S rRNA gene amplicon sequencing

16S rRNA gene amplicon sequencing was performed using the Illumina MiSeq platform and customized sequencing primers at Argonne National Laboratory. Briefly, DNA was extracted from fecal samples using the QIAamp Fast DNA Mini Stool Kit and quantified using a NanoDrop 1000 Spectrophotometer. Extracted DNA was sent to Argonne National Laboratory where the V4 region of the 16S rRNA gene was amplified from each DNA sample using the modified primers 515F: GTGYCAGCMGCCGCGGTAA and 806R: GGACTACNVGGGTWTCTAAT (32, 33). A unique forward primer containing 12 bp barcode was used to facilitate multiplexing of the samples (34). Next, cleaned PCR products were quantified and pooled at equimolar concentrations for paired-end sequencing (151 bp by 12 bp by 151 bp) on the Illumina MiSeq platform.

### 16S rRNA gene amplicon data processing

Raw 16S rRNA gene amplicon sequences were imported into QIIME 2 (version 2018.6) (35) and demultiplexed using the sample-specific barcodes. Demultiplexed reads were denoised and dereplicated to obtain an amplicon sequence variants (ASVs) table using ‘dada2-denoise-paired’ plugin. The ASV table was rarified at 8,384 sequences per sample and ASVs were classified to the lowest possible taxonomic rank using QIIME’s feature-classifier plugin and the Silva 132 99% OTUs full-length sequences and was purged of ASVs assigned to mitochondria, chloroplasts, or were unidentified at the domain level. The resulting filtered ASV table with the taxonomic assignment, ASV phylogenetic tree, and associated metadata were imported in R (3.5.0) (https://www.R-project.org/) for further analyses using phyloseq (version 1.25.2) (36) and vegan (version 2.5.2).

### Microbial diversity and statistical analyses

To estimate alpha diversity, Shannon, Simpson, and Observed ASVs diversity indices were calculated. To test if there were significant differences in alpha diversity in response to diet, antibiotic administration, and CDI, a two-way mixed ANOVA and a Bonferroni test were performed to conduct comparisons of alpha diversity indices with correction for multiple comparisons in SPSS (version 25). A few data points are missing data due to hamster death. The resulting analysis was visualized using ggplot2 (version 3.0.0) and modified in Inkscape. NMDS was performed based on Bray-Curtis dissimilarity to examine relationships between the samples with respect to diet, antibiotic treatment, and CDI over the course of experiment, and hypotheses were tested using ANOSIM. Further, similarity percent (SIMPER) analysis was performed to identify ASVs that contributed to 50% of the observed differences in microbial communities during the study (37).

One animal, hamster #9 in the Standard Lab Diet group was shown to be a statistical outlier in nearly all analyses since it showed no apparent microbiome disruption (e.g., alpha diversity, NMDS, *C. difficile* relative and absolute abundance) and was removed from figures and statistical calculations. Similar host-specific variability has been observed in diet-microbiome studies (38, 39) and may be related to rejection of the clindamycin gavage in our study.

### Absolute quantification of *C. difficile*

Primers targeting a subunit of the *C. difficile* D-proline reductase proprotein *prdA* (primers oLB170/oLB171) reported in Bouillaut et al. 2013 (40) were used for absolute quantification of *C. difficile* in fecal DNA on key days from the experimental timeline (**Figure 1**) (Days 2, 3, 8, and 11), using a *C. difficile* R20291 *prdA* PCR product as a standard. Briefly, triplicate qPCR reactions were set up (10 μL each) containing 1 μM of *prdA*-specific forward and reverse primers, metagenomic DNA as a template, and PerfeCTa SYBR Green SuperMix (Quanta Biosciences, Gaithersburg, MD, USA). The qPCR assays were run on CFX96 Touch™ Real‐Time PCR Detection System (Bio‐Rad, Hercules, CA, USA) using the following PCR conditions: initial denaturation at 95 °C for 3 mins, followed by 40 amplification cycles at 95 °C for 10 sec, 53 °C for 10 sec and 72 °C for 15 sec (40). A standard curve was generated from serial dilutions of a known concentration of *prdA* PCR products within each qPCR assay, and triplicate counts of *prdA* were averaged to estimate *prdA* copy number for each metagenomic sample. Additionally, a melt curve analysis was performed to check amplification specificity. For all assays, PCR efficiency was above 90% with a correlation coefficient > 0.99.

## Results

### The high-carbohydrate diet exaggerated antibiotic- and CDI-associated microbiome diversity loss

Over 13.9 million high-quality 16S rRNA gene fragment reads were obtained from 447 fecal samples collected along the experimental timeline (**Figure 1**), with a mean of 30,085 reads per sample, representing 3,533 unique ASVs. After rarefying the ASV table to 8,384 sequences per samples, a total of 410 samples were retained. To understand changes in the gut microbiome due to experimental diets within the context of the CDI time course, alpha diversity was analyzed using observed amplicon sequence variants (ASVs) (**Figure 2**), Simpson’s evenness (**Figure S1**), and Shannon diversity (**Figure S2**). Untreated, uninfected control hamsters fed the standard lab diet (Standard Lab Diet (-CDI) group) showed no significant change in alpha-diversity metrics over time (p > 0.05, two-way mixed ANOVA). In contrast, there were significant changes for all three diversity metrics over time (p < 0.05, two-way mixed ANOVA) for both experimental groups. In the standard lab diet group, this constituted a >90% reduction of observed ASVs during antibiotic treatments and CDI compared to pre-antibiotic levels (**Figure 2**). For these hamsters, ASVs richness increased in surviving animals by the end of the experiment, but was still well below a pre-antibiotic, pre-infection state at the end of the experiment (**Figure 2**, Day −1 mean 282.8 ±34.51 S.D. versus Day 14 mean 98.5 ±34.0 S.D.). This effect was exaggerated for animals in the high-carbohydrate diet group, where mean community richness was depleted by >98% by Day 2 (**Figure 2**, Day −1 mean 205.2 ±21.7 S.D. versus Day 2 mean 19.9.0 ±5.9 S.D., p < 0.05) and did not rebound for the duration of the experiment. There were also significant daily decreases in diversity in the high-carbohydrate diet group microbiomes after clindamycin administration (Day −1 versus Day 0), after the first dose of vancomycin and *C. difficile* challenge (Day 0 versus Day 1), after the second dose of vancomycin (Day 1 versus Day 2), and after the final dose of vancomycin (Day 2 versus Day 3) (p < 0.05, two-way repeated-measures ANOVA).

**Fig 2.**
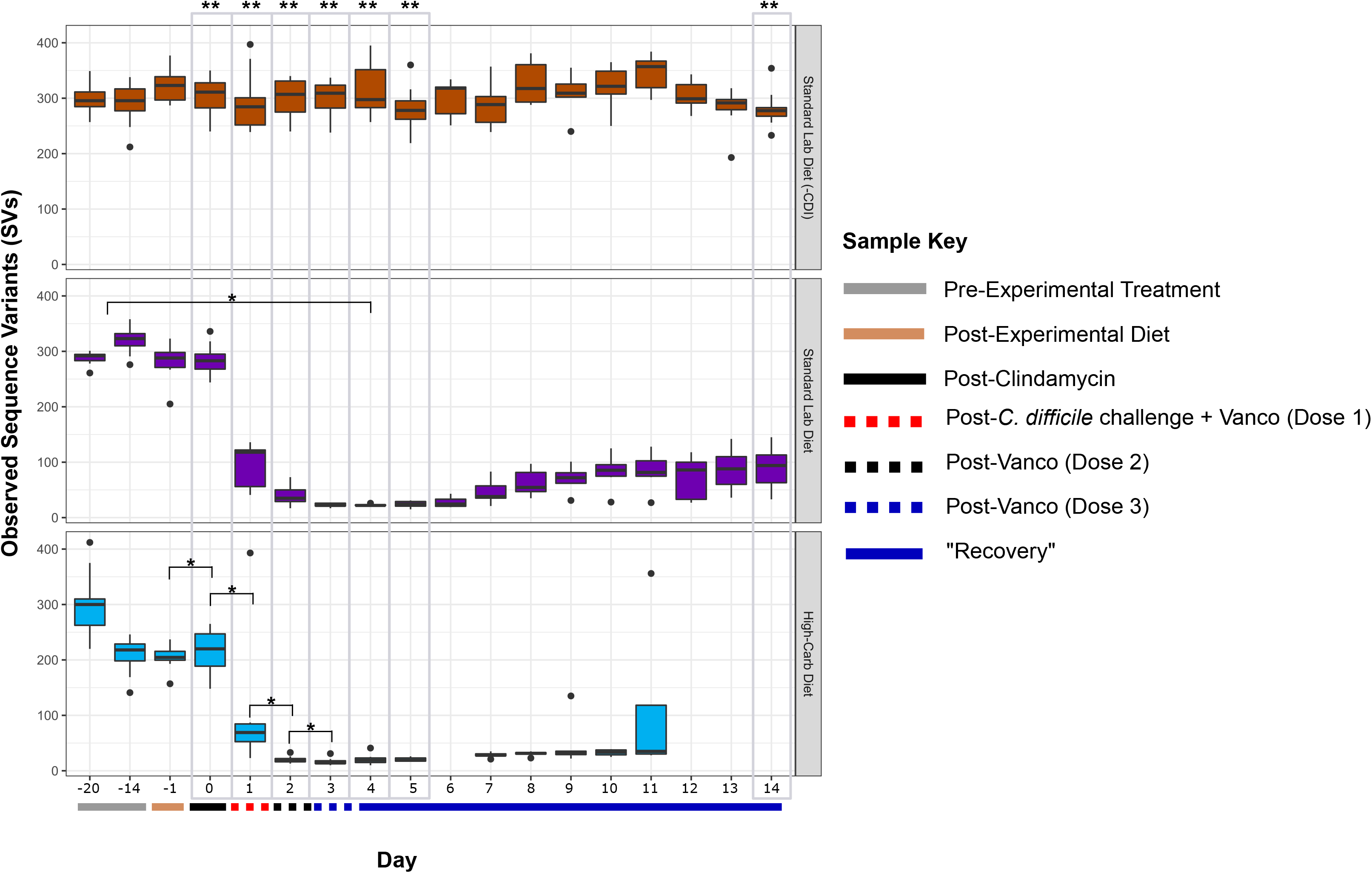
Effect of diet on microbial richness. Observed sequence variants (SVs) is shown for uninfected hamsters fed a standard lab diet (orange, n = 10), and infected hamsters fed a standard lab diet (purple, n = 9), or high-carbohydrate diet (blue, n = 10). Administration of experimental diets (solid tan line, x-axis), timepoints after antibiotics (dashed lines, x-axis) and *C. difficile* challenge (black line, x-axis) are indicated. Black dots above and below boxplots represent outliers. (*) Indicates significant (p < 0.05) loss of diversity in within-group pairwise comparisons shown by the brackets. Gray boxes highlight comparisons between groups after a change in diet on Day −1, antibiotic treatments on Days 0-3, post-infection on Days 4 and 5, and recovery on Days 14. (**) indicate significant (p < 0.05) difference between groups on a given day (repeated measures mixed ANOVA).

### Antibiotics and CDI disrupted microbial community composition, which was prolonged in animals fed the high-carbohydrate diet

To further assess microbial community alterations over time, Bray-Curtis dissimilarity was visualized by nonmetric multidimensional scaling (NMDS) (**Figure 3, Figure S3**). Pre-antibiotic diet-specific communities did not emerge, as indicated by overlapping “Pre-Experimental” (gray solid line) and “Post-Experimental Diet” (tan solid line) confidence ellipses (ANOSIM R = 0.15, p-value = 0.18). In the experimental groups, the microbial communities were disrupted by antibiotic treatments and CDI, as indicated by progressive shifts in ellipses associated with the three-day vancomycin regimen and *C. difficile* spore challenge (red, black, and blue dashed ellipses). Moreover, the vancomycin treatment- and CDI-associated ellipses were distinct between hamsters based on diet, demonstrating a synergistic interaction between diet and antibiotic treatment/CDI that changed the gut microbiome composition through the full course of the experiment: post-*C. difficile* challenge and vancomycin dose 1 (red ellipses, ANOSIM: R = 0.5, p= 0.001); post-vancomycin dose 2 (black ellipses, ANOSIM: R = 0.64, p = 0.001); post-vancomycin dose 3 (blue ellipses, ANOSIM: R = 0.89, p = 0.001); and recovery phase (blue ellipses, ANOSIM: R = 0.57, p = 0.001). Full microbiome recovery was not seen in surviving animals in either experimental group, though the microbiomes in the standard lab diet group approached the pre-experimental condition, whereas those in the high-carbohydrate diet group did not change beyond the acute phase of CDI (blue ellipses). The microbiomes in the negative control group (Standard Lab Diet (-CDI)) did not vary throughout the experiment (**Figure S4**).

**Fig 3.**
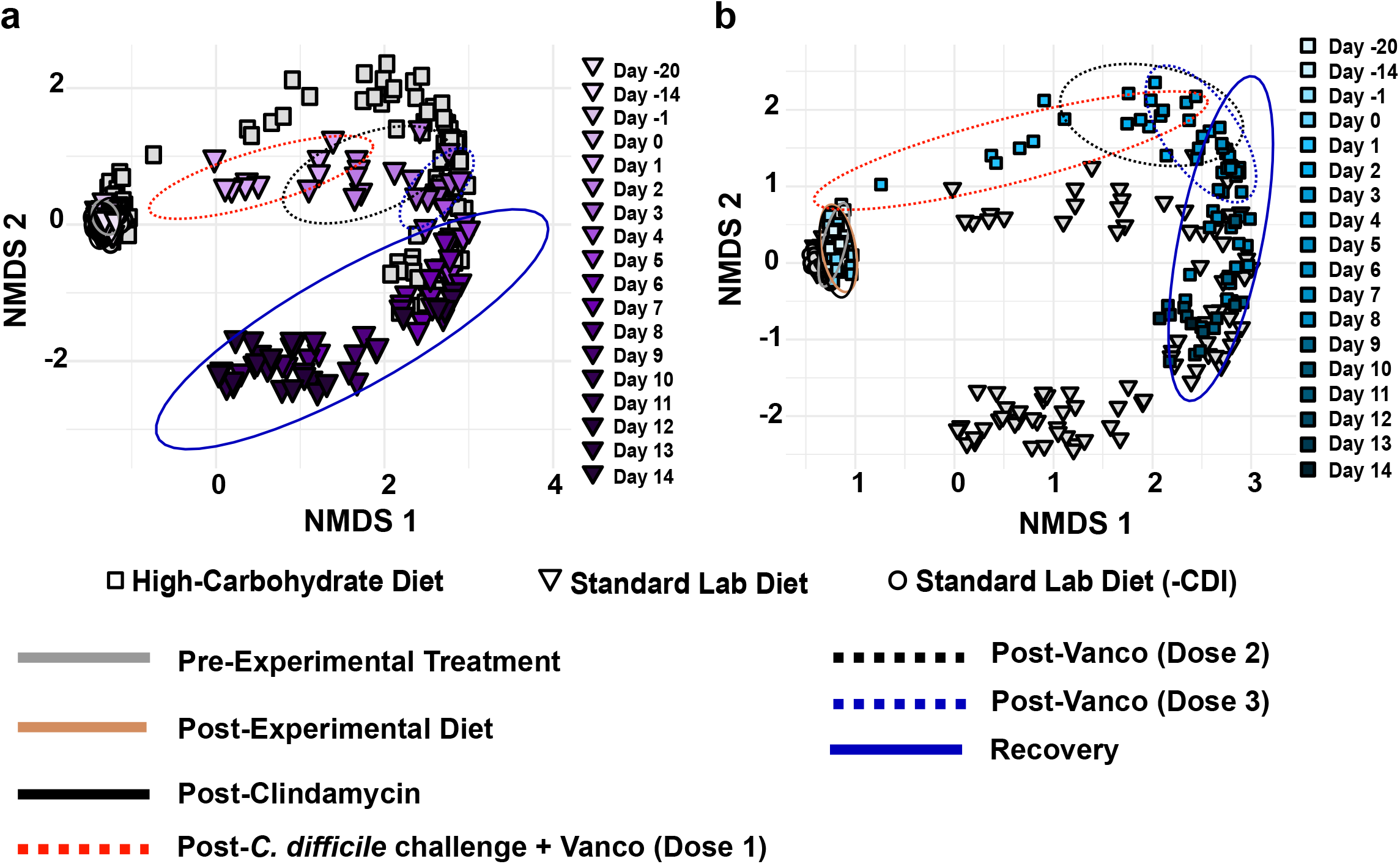
NMDS analysis based on Bray-Curtis dissimilarity. Each panel is a visualization of the same data and highlights the analysis for infected hamsters fed a) the standard lab diet (purple, triangles) and b) the high-carbohydrate diet (blue, squares). Colors are shaded to show time progression through the experiment. Ellipses represent standard errors of the mean (95% confidence) for samples associated with the standard lab diet (Pre-Experimental Treatment, gray solid line, Days −20, −14, −1), the high-carbohydrate microbiome (Post-Experimental Treatment, tan solid line, Day −1), the effect of clindamycin (Post-Clindamycin, black solid line, Day 0), the first dose of vancomycin and CDI challenge (Post-*C. difficile* challenge + Vanco (Dose 1), red dashed line, Day 1), the second dose of vancomycin (Post-Vanco (Dose 2), black dashed line, day 2), the third dose of vancomycin (Post-Vanco (Dose 3), blue dashed line, day 3), and the recovery phase (solid blue line, days 4-14).

These changes in beta diversity were consistent with changes in phylum-level relative abundance, which showed dramatic shifts in the microbiomes for both experimental groups beginning on Day 1. These changes are characterized by large expansions of *Proteobacteria* in both groups, which coincided with antibiotic treatment and acute CDI. The high abundance of *Proteobacteria* largely subsided by Day 4 in the standard lab diet group microbiomes (**Figure S5**) but persisted in the high-carbohydrate diet group microbiomes (**Figure S6**). There were smaller expansions of *Epsilonproteobacterota* (specifically *Campylobacteraceae*) coincident with antibiotic treatment and acute CDI in both experimental groups, which similarly persisted longer in the high-carbohydrate diet group microbiomes (**Figure S6**). Only the standard lab diet group microbiomes showed a transient increase in *Tenericutes* during CDI, which quickly decreased to pre-experimental conditions. In contrast, *Tenericutes* became depleted during antibiotic treatment and acute CDI in the high-carbohydrate diet group microbiomes and never recovered. *Fibrobacteretes* became depleted during antibiotic treatment and acute CDI in both experimental groups and also never recovered. Some animals in both experimental groups showed increases in *Fusobacteria* during recovery from acute CDI, and some animals in the high-carbohydrate diet group showed increases in *Actinobacteria*. The microbiomes in the negative control group (Standard Lab Diet (-CDI)) did not vary throughout the experiment (**Figure S5**).

### Diet-specific microbiome changes were associated with antibiotic treatments and CDI and recovery

SIMPER analysis was performed to identify ASVs that contributed most to the observed changes in beta diversity (**Figure 4**). This analysis identified 44 ASVs that cumulatively accounted for 30% of microbial community dissimilarity between all pairwise comparisons of the three animal groups throughout the experiment. Almost one-third of these ASVs (13 ASVs) belonged to the family *Muribaculaceae*. Nearly all *Muribaculaceae* ASVs decreased in abundance after administration of the first dose of vancomycin and *C. difficile* spores, with recovery of one of the *Muribaculaceae* ASVs (*Muribaculaceae.*0) and emergence of a new *Muribaculaceae* ASV (*Muribaculaceae.*2) only in hamsters in the Standard Lab Diet group late in the experiment.

**Fig 4.**
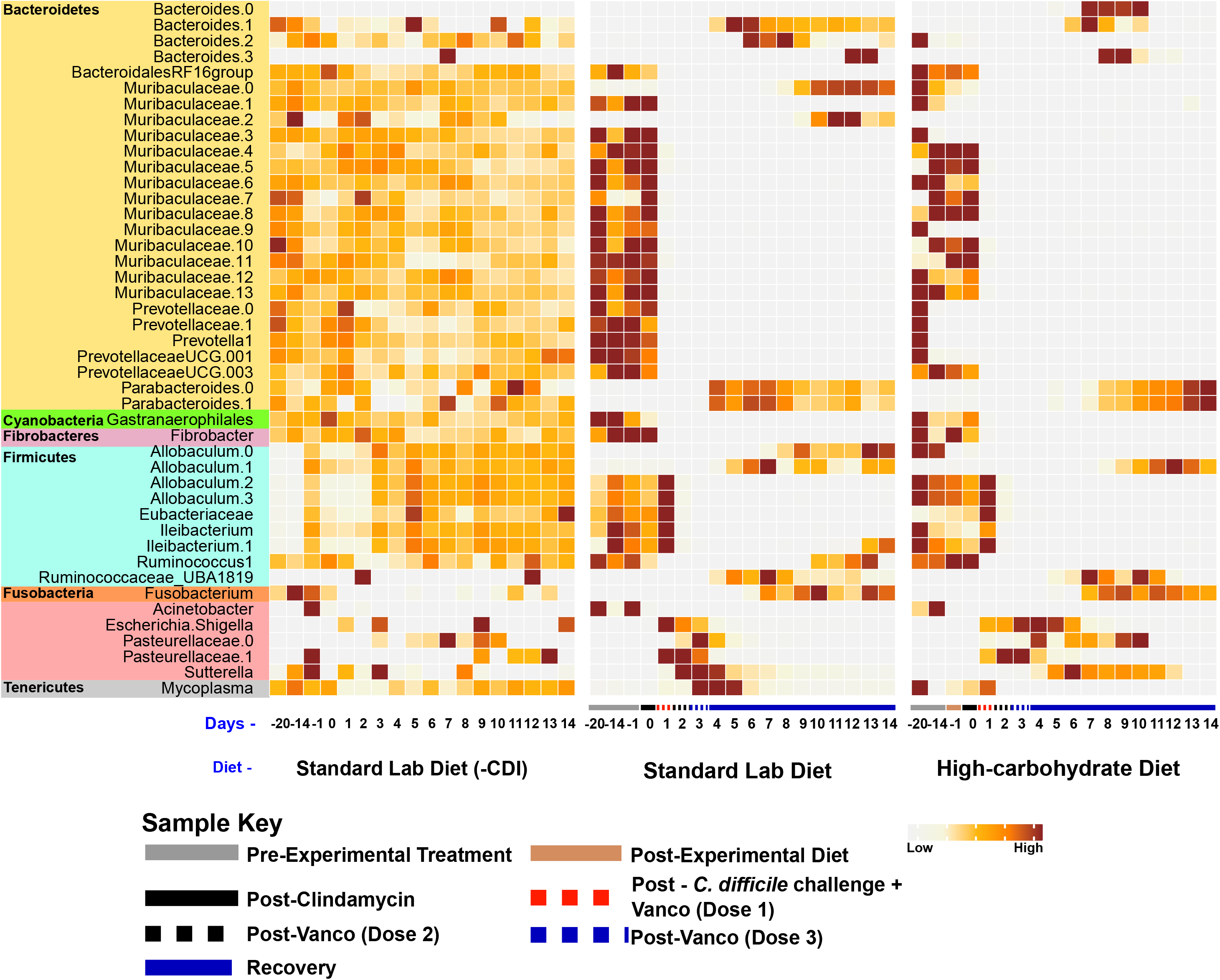
Heatmap of 124 ASVs identified using SIMPER analysis that contributed to 50% of observed dissimilarity. Results display top SVs responsible for dissimilarity between experimental groups. The heat map indicates the mean relative abundances of 124 SVs that contributed cumulatively to 50% of community dissimilarities at each time point among the hamsters fed the standard laboratory diet and high-carbohydrate diet. Each square represents the mean relative abundance of the given SV on a particular day for a particular diet. Higher intensity of brown coloring correlates with higher relative abundance. Asterisks (*) indicate taxa featured in the manuscript text.

Several other taxa also decreased following antibiotics and CDI in both experimental groups, with some showing diet-specific responses. For example, one *Ileibacterium* ASV (*Ileibacterium.1*) and *Ruminococcus* recovered only in the Standard Lab Diet group. Several *Prevotellaceae* ASVs were similarly depleted following antibiotic treatment and CDI in the standard lab diet group, but they were already depleted following the change in diet in the high-carbohydrate group before antibiotic treatment. None of these *Prevotellaceae* ASVs recovered during the experiment. Similarly, an unassigned member of the *Gastranerophilales* within the *Cyanobacteria*, a member of the genus *Fibrobacter* within the *Fibrobacteres,* and several ASVs assigned to the *Firmicutes* (i.e., *Eubacteriaceae* (1 ASV), one ASV belonging to *Ileibacterium* (2 ASVs), and an unassigned *Eubacteriaceae* ASV (1 ASV)) were depleted due to antibiotics and CDI and never recovered.

Other groups increased in abundance in response to dysbiosis. There was a transient increase after antibiotic administration in members of the phylum *Proteobacteria*, which included ASVs assigned to *Acinetobacter* (1 ASV), *Escherichia*/*Shigella* (1 ASV), *Pasteurellaceae* (2 ASVs), and *Sutterella* (1 ASV). Outside the *Proteobacteria*, some other taxa showed similar patterns, including *Parabacteroides* (2 ASVs), *Fusobacterium* (1 ASV), and *Mycoplasma* (1 ASV). Four members of the genus *Bacteroides* also expanded during dysbiosis but showed diet-specific patterns. *Bacteroides.0* expanded only in the high-carbohydrate diet group, whereas *Bacteroides.2* expanded only in the standard lab diet group. The two other *Bacteroides* ASVs expanded in both experimental groups.

### The high-carbohydrate diet prolonged *C. difficile* carriage

Relative quantification of *C. difficile* prolonged carriage in the high-carbohydrate diet group. The relative abundance of *C. difficile* among ASVs was similar in the two experimental groups on Day 2 (2 days post-infection) (**Figure 5**). However, *C. difficile* dropped below the detection limit for all surviving hamsters in the standard lab diet microbiomes by Day 3 and remained undetectable for the remainder of the experiment. In contrast, *C. difficile* remained detectable in surviving hamsters in the high-carbohydrate diet group from Day 2 to Day 10.

**Fig 5.**
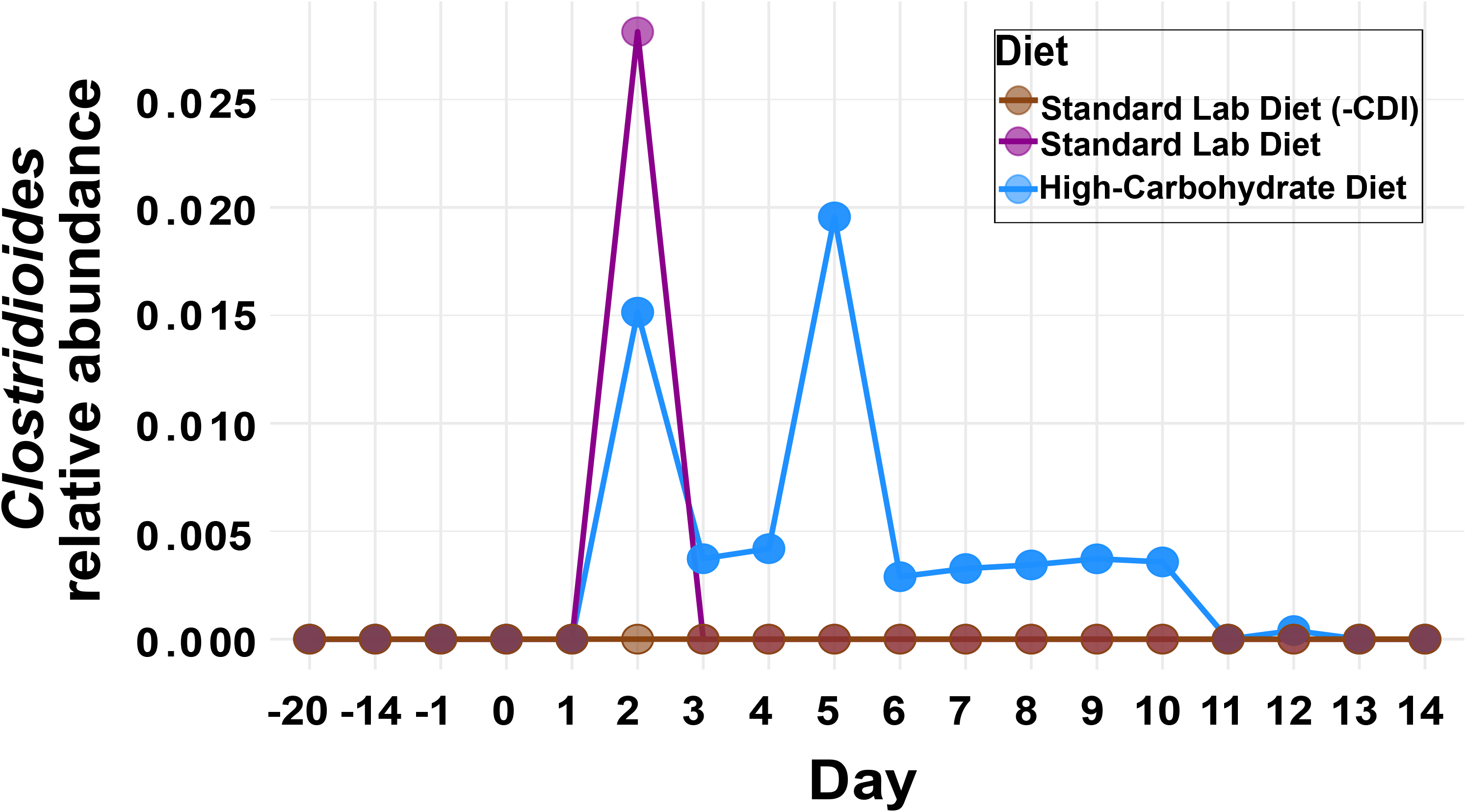
*C. difficile* relative quantification. a) 16S rRNA amplicon-based relative abundance of *C. difficile* across the experimental timeline. Each dot represents the mean relative abundance of *C. difficile* on a particular day for a given diet.

### The high-carbohydrate diet increased secondary CDI and mortality in hamsters

Hamster mortality was documented to assess the effect of the high-carbohydrate diet on hamster survival after treatment with sub-clinical concentrations of vancomycin and challenge with *C. difficile* spores on Day 0 until the end of the experiment on Day 14 (**Figure 6**). Three independent statistical tests were run to assess mortality. Although differences in the full experimental course or the acute phase of CDI (Days 1-4) were not statistically significant (p > 0.05, log-rank test), there was a significant difference between mortality in the later phase of CDI (Days 5-14; p = 0.051, log-rank test), documenting a higher incidence of mortality due to delayed-onset CDI in animals fed the high-carbohydrate diet. Three of the nine hamsters fed the Standard Lab Diet and infected with *C. difficile* developed severe CDI and were euthanized after 48 hours of infection (33% mortality) during the acute phase of CDI, while the remaining six hamsters survived for the entirety of the experiment. In comparison, two of the ten hamsters fed the high-carbohydrate diet and infected with *C. difficile* developed severe CDI during the acute phase and were euthanized after 48 hours of infection; one more hamster died within the next 24 hours, after which no mortality was observed until Day 8. Five additional hamsters fed the high-carbohydrate diet died on Day 9 (2 hamsters), Day 10 (1 hamster), and Day 12 (two hamsters), leading to a total of 80% mortality of the hamsters fed the high-carbohydrate diet. All control hamsters fed the standard lab diet but not infected (Standard Lab Diet (-CDI)) survived for the duration of the experiment.

**Fig 6.**
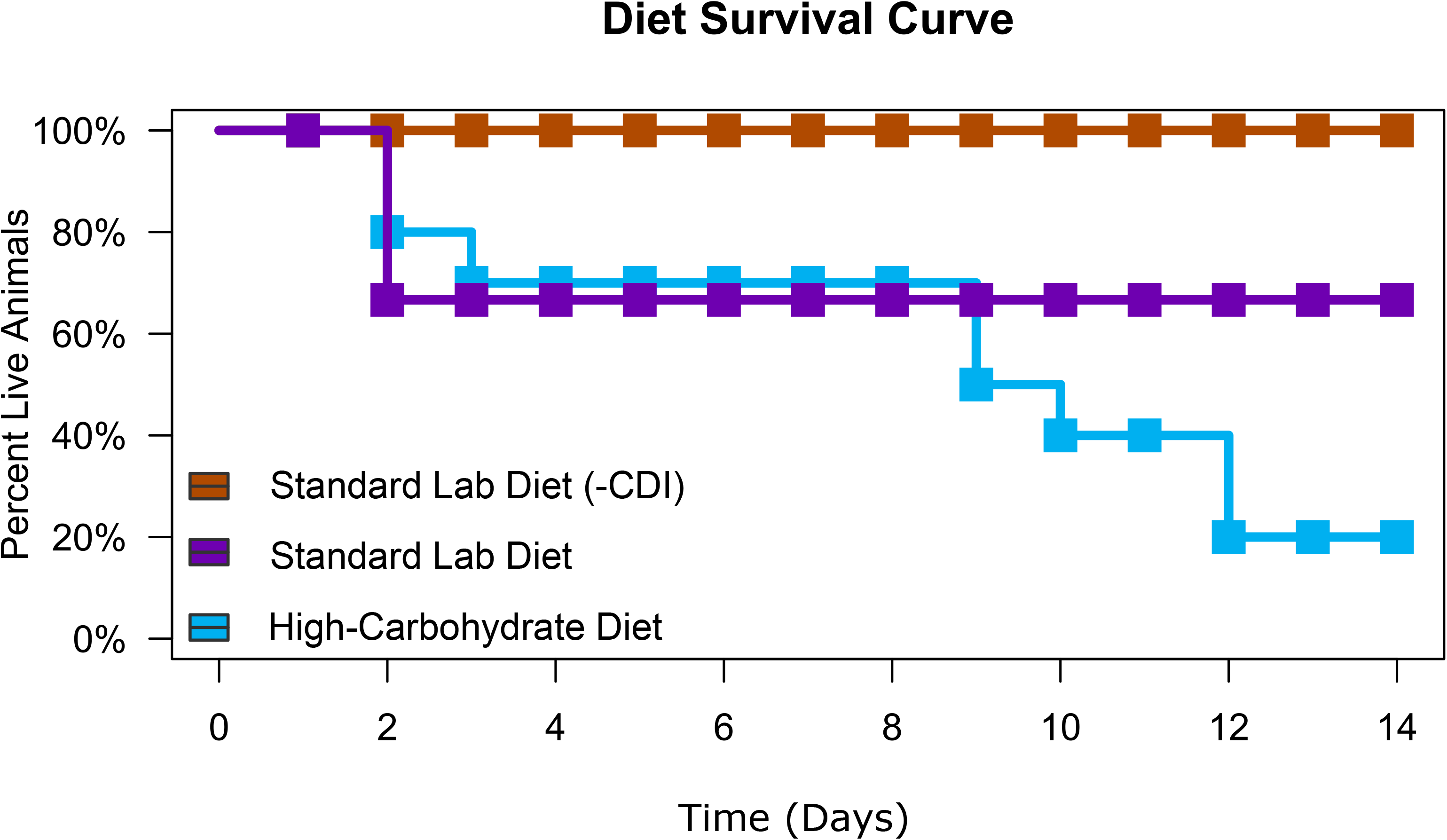
Effect of diet on hamster survival. Kaplan-Meier survival curves for uninfected hamsters fed the standard lab diet (orange, n = 10), and infected hamsters fed the standard lab diet (purple, n = 9) and high-carbohydrate diet (blue, n = 10). There were no significant reductions in survival (p > 0.05, log-rank test) between the two experimental groups for the full experiment or for the acute phase of CDI (Days 0-4); however, there was a significantly higher mortality in the High-Carbohydrate Group later in the experiment due to delayed-onset CDI (Days 5-15; p = 0.01, log-rank test).

## Discussion

Syrian golden hamsters have been instrumental as experimental models to understand various aspects of *C. difficile* biology and pathogenesis; however, the effects of diet on CDI in hamster models of disease have not been fully explored. Here, a standard lab diet and a high-carbohydrate diet containing simple sugars were directly compared in an antibiotic-associated CDI model in hamsters to explore their effects on CDI development and microbial community dynamics.

### Effect of antibiotics, CDI, and diet on gut microbiomes in hamsters

Conventionally, reduced gut microbial diversity is considered as a hallmark of CDI susceptibility and the loss of colonization resistance (41, 42). However, more recent reports indicate CDI susceptibility is not specifically related to microbial diversity and suggest that diet plays a more important role in CDI outcome (17, 18). For example, the microbial-accessible carbohydrate inulin was shown to increase short-chain fatty acid production, reduce *C. difficile* burden, and promote CDI resolution in a mouse model of disease, with no significant effect on microbiome diversity (17). More strikingly, in our previous study using mice, we showed that a high-carbohydrate diet rich in sucrose and digestible starch dramatically reduced microbiome richness and diversity during antibiotic treatment and CDI compared to mice on the standard lab diet, yet the high-carbohydrate diet prevented CDI signs in almost all animals and those that showed mild CDI signs quickly recovered. Here we observed a similar effect of the high-carbohydrate diet on microbial diversity in hamsters (**Figure 2; Figure 3**); however, in this case lower microbial diversity correlated with poor long-term prognosis (**Figure 6**).

Despite overall similar effects of this high-carbohydrate diet on the microbiomes in mouse and hamster experimental models, the dynamics of these microbial communities were distinct. For example, mice developed distinct microbiomes in response to the high-carbohydrate diet, before antibiotic treatment and *C. difficile* challenge. That change was related to depletion of several members of the *Lachnospiraceae* and *Ruminococcaceae*. Although pre-CDI, diet-specific microbiomes were not evident in hamsters (**Figure 3**), a depletion of some *Prevotellaceae* was observed following the change in diet (**Figure 4**). In both models and both diets, antibiotic treatment and CDI was accompanied by a proliferation of *Proteobacteria*. Many gut *Proteobacteria*, particularly *Enterobacteriaceae*, overgrow in response to inflammation, due to respiration of reactive nitrogen species-derived nitrate and nitrite (by denitrification and dissimilatory nitrite reduction to ammonium), which promotes a pro-inflammatory positive feedback loop (43). In this study, a more diverse group of respiratory *Proteobacteria* expanded during dysbiosis and CDI (compared to results in mice), which may be related to the more sensitive nature of hamsters to CDI and/or the effects of vancomycin, which was not used in the mouse CDI model. Blooms of *Proteobacteria* in the dysbiotic gut have been documented in many microbiome studies performed in humans and mice with CDI (44, 45).

A very striking response to antibiotics and CDI in hamsters was the depletion of many members of the *Muribaculaceae*, making up almost 30% of ASVs that significantly changed through the course of the experiment (**Figure 4**). Although very few members of the *Muribaculaceae* have been cultivated in the laboratory, both metagenomics and cultivation studies show that they have an abundance of glycoside hydrolases and ferment MACs to SCFAs (46–50). The fact that many members of the *Muribaculaceae* were greatly reduced during CDI in hamsters suggests a loss of SCFA production, which has been associated with decreased gut health due to changes in gut epithelia morphology and permeability, and poor CDI outcomes (17, 51–53). However, the expansion of some other *Muribaculaceae* during and after CDI in the standard lab diet group complicates this interpretation and begs for more detailed study of individual species and strains in the *Muribaculaceae*. There was also a decrease in relative abundance of members of the phyla *Cyanobacteria*, *Fibrobacteres*, *Firmicutes*, and *Tenericutes* after administration of antibiotics regardless of diet (**Figure 4; Figures S5, S6**). This loss in specific taxa due to antibiotic treatment is consistent with previous research in mice (42, 54).

The microbiome was somewhat restored by the end of the experiment (**Figure 3**), but only in the standard lab diet microbiomes, as evidenced by recovery of some ASVs, including some members of the genera *Bacteroides*, *Ruminococcus*, *Ileibacterium,* and *Muribaculaceae* (**Figure 4**). The appearance of these ASVs during the recovery phase was also reflected NMDS plots based on Bray-Curtis dissimilarity showing the Day 14 microbiomes approaching the “Pre-Experimental Treatment” time points for Standard Lab Diet microbiomes (**Figure 3; Figures S5, S6**). This provides evidence of partial restoration of the gut microbial community after antibiotic-induced CDI, which is consistent with studies that examine the effect diet on CDI outcomes in mice.

### Primary, acute antibiotic-associated CDI may be independent of diet in hamsters yet mortality due to delayed-onset CDI was not

Primary, acute antibiotic-associated CDI appeared to be independent of diet in this experiment, as evidenced by mortality of three animals from each group during the acute phase (within 72 hours of *C. difficile* challenge (**Figure 6**) and the detection of *C. difficile* in the 16S rRNA gene amplicon data (**Figure 5**). Although not significant, the lower relative and absolute abundance of *C. difficile* in the High-Carbohydrate group was generally consistent with the known role of carbohydrates to mitigate primary CDI. For example, a previous study in which hamsters were fed a liquid diet supplemented with 1.4% soy fiber showed lower *C. difficile* toxin positivity and stool liquidity compared to control animals during the acute phase of CDI (30). In that study, the milder acute CDI led to a 34% increase in survival time, although all hamsters succumbed to acute CDI within 7 days post-infection. Similar improvements in mean survival time were observed in another study in which hamsters were fed fructooligosaccharides (30 g/L drinking water) (29); however, the protective effect of fructooligosaccharides was not observed in *C. difficile*-infected hamsters that were treated with vancomycin.

### Simple sugars promote *C. difficile* carriage in animal models and may later become symptomatic

Despite similar CDI courses during the acute phase (within 72 hours of infection), the high-carbohydrate diet led to a poor prognosis, particularly due to delayed-onset CDI later in the experiment (**Figure 6**). To our knowledge, dietary carbohydrates have not previously been shown to exacerbate CDI in any animal models. We propose that this unusual result was a product of prolonged dysbiosis and *C. difficile* carriage due to synergistic effects between antibiotics, the high-carbohydrate diet, and the extreme sensitivity of hamsters to CDI.

In this study, *C. difficile* was detectable in the high-carbohydrate diet group microbiomes by Day 2 and remained so for most of the experiment (**Figure 5**). Strikingly, *C. difficile* was not detected in the Standard Lab Diet amplicon dataset after Day 2, despite the deep sequencing effort. The persistence of *C. difficile* in the high-carbohydrate diet microbiomes (≥7×10^4^ gene copies per feces for Days 2, 3, 8, and 11) may be related to the greater extent and duration of dysbiosis in these animals. Microbiome richness was severely depleted (>98% reduction in ASVs (**Figure 3**)) by Day 2 in these animals and did not increase significantly for the duration of the experiment. Hamsters in the high-carbohydrate diet suffered from a second period of mortality triggered days after a second *C. difficile* bloom was detected. Similarly, Bray-Curtis analyses showed that high-carbohydrate diet microbiomes remained dysbiotic throughout the experiment (**Figure 3**) and many ASVs did not recover (**Figure 4**). Very similar microbiome changes were seen previously using these exact same diets in mice (18), where *C. difficile* persisted in animals fed the high-carbohydrate diet but appeared to be cleared from mice on the standard lab diet by the end of the experiment, 30 days after *C. difficile* challenge. Similarly, final microbiome richness was >2x higher for mice fed the standard lab diet compared to the high-carbohydrate diet and the high-carbohydrate diet microbiome was never fully restored (**Figure 2**).

Despite similar microbiome responses, the outcomes of CDI in hamsters and mice on these two diets were opposite. We propose that these different outcomes relate to the tolerance of these two animals to *C. difficile*. Although hamster models of CDI mirror the classic features of human CDI including the shifts in gut microbial communities, diarrhea and histological anomalies, and more advanced changes such as development of pseudomembranes, it is very different from murine models of CDI. Hamsters are highly susceptible to CDI; they show severe and sudden onset of CDI symptoms with as little as a few dozen spores (55, 56) and require suboptimal doses of vancomycin to prevent *C. difficile* overgrowth and death (56, 57). In contrast, mice are resistant to CDI development and require a greater spore load (~10^8^ spores), an aggressive antibiotic regimen, and are able to recover from infection within a few days (22, 58–61). Mice models are also slow to develop CDI and display a wide range of CDI signs (22, 62). We therefore speculate that inherent differences between host susceptibility to CDI led to the observed differences in the CDI outcome (18). In the case of the hamster experiments described here, we propose that vancomycin administered to animals on Days 0-3 limited *C. difficile* proliferation till the vancomycin was cleared from the system, after which *C. difficile* proliferated and led to severe, acute CDI, and death beginning on Day 8.

## Conclusions

This study, along with our recent study in a mouse CDI model, documented persistent dysbiosis in animals fed a high-carbohydrate diet following antibiotic treatment and CDI. The dysbiotic, high-carbohydrate diet microbiome was conducive to prolonged *C. difficile* carriage in both experimental systems. While mice on the high-carbohydrate diet were protected from CDI and persisted in an asymptomatic carrier state, *C. difficile* proliferated in hamsters following termination of vancomycin treatment, leading to higher mortality in the high-carbohydrate diet group. The divergent long-term prognoses of the two experimental models in response to the high-carbohydrate diet may translate to different diet-specific responses in humans. Additionally, if high-carbohydrate diets promote asymptomatic *C. difficile* carriage in humans, it is possible that changes in diet, host health, antibiotics, or other perturbations may lead to *C. difficile* outgrowth and expression of CDI. Thus, we advocate a careful, context-dependent interpretation of results when extrapolating the findings from animal models studies to clinical use.

## Data Availability

Files containing the original unfiltered sequences are available from the NCBI-SRA under accession number PRJNA766033.

## Acknowledgements

This research was supported by the NIH under grant 1R01AI109139-01A1, an INBRE pilot grant under NIH GM103340, and a UNLV Faculty Opportunity Award. *C. difficile* R20291 (RT027) was a gift from Nigel Minton, University of Nottingham.

